# Passive Physiological Responses Fail to Predict Context-Dependent Action Selection in Bats

**DOI:** 10.64898/2026.06.09.731189

**Authors:** Zaria George, Inaky Marin, Anna C. Sanderson, Angeles Salles

## Abstract

How social animals encode vocalizations, assign them value, and formulate behavioral responses remains largely unknown. We asked whether physiological signatures of social-call perception predict behavioral responses across contexts, using the Egyptian Fruit Bat (*Rousettus aegyptiacus*), an auditory specialist with a rich repertoire of social calls. Using heart rate monitoring during playback of conspecific vocalizations, we found that females showed larger heart rate responses than males to social calls (aggression and distress), whereas non-social echolocation calls evoked no sex difference. In vivo recordings from primary auditory cortex (A1) revealed call-selective units whose selectivity was independent of frequency tuning, with the largest selective fraction for distress calls. In a behavioral assay, female bats approached a distress-call playback only when a live conspecific was coupled with it. However, in isolation, the same calls elicited interest (grooming, pointing) but no approach. Together, the neural and autonomic signatures of social-call perception are present across contexts, whereas the behavioral response is not. Social context, therefore, does not modulate the behavioral readout of social calls; rather, it gates it.

**Significance Statement:** How a social animal converts the perception of a vocalization into behavior remains poorly understood. Using the Egyptian Fruit Bat (*Rousettus aegyptiacus*), we show that physiological and neural responses to social calls are present in isolation, whereas approach behavior is not. We conclude that social context acts as a necessary gate between sensory representation and action, not a modulator that adjusts an existing response.

## Introduction

Social communication is fundamental to our daily interactions and largely depends on promptly and accurately interpreting auditory information. For this communication to be successful, the appropriate filtering of sensory information is critical for salient awareness and the recognition of social cues. By examining how social vocalizations are encoded across contexts, we can better identify where in the auditory pathway value is assigned to these calls and whether context influences perception. Studies have shown that the primary auditory cortex (A1) combines the linear, tonotopic representation of frequency with a nonlinear representation, allowing the system to become sensitive to spectral features independent of a specific MUA’s best frequency [1-3]. The integration of auditory information occurring in A1 makes it an ideal hub for investigating social sound processing. Several studies have investigated these processes in rodent models, examining the representation of Ultrasonic Vocalizations (USVs) in A1 [4,5], including maternal neuronal responses to pup isolation calls [6] and physiological responses to USV playbacks in mice [7]. In gerbils, the auditory cortex has been found to encode the categorization of a conspecific vocalization and its identity based on acoustic features [8]. Although this work provides valuable insight into the auditory processing of social sounds across taxa, a more comprehensive characterization requires a combined physiological and behavioral investigation in a species with a richer call repertoire to directly examine the role of social context.

The Egyptian Fruit Bat, *Rousettus aegyptiacus*, emerges as a strong model for this study, as its repertoire contains multilayered information that conveys the caller’s identity and emotional state, as well as the outcome of social interactions [9]. These bats utilize social calls to navigate environments where social dynamics constantly change. In this species, mating, isolation, distress, and aggression calls have been classified according to the behavioral context in which they were emitted [10]. How such calls are encoded and how they shape behavior across contexts remains unknown. In the Egyptian Fruit Bat, we measure physiological responses to conspecific call types, then test whether these signals are sufficient to drive behavior.

## Results

### Heart Rate Responses During the Perception of *R. aegyptiacus* Vocalizations are Sex-Dependent

Heart rate recordings were obtained from 12 bats (6 females, 6 males) across three consecutive days of session recordings. When the dataset was divided by sex, differences emerged during playback of socially relevant vocalizations, specifically distress and aggression (Mann-Whitney U-test, p < 0.001). This was reflected in the median change in heart rate per sex across groups (**Figure 2a**) and in the pattern of heart rate change during stimulus presentations (**Figure 2b**). To test this statistically, a Mann-Whitney U test revealed that males and females responded differently across all stimuli (**p < 0.001; Figure 2c**). A Kruskal-Wallis test followed by Dunn’s post-hoc test identified the social stimuli (distress and aggression) as the primary drivers of this difference (**p < 0.05; Figure 2d**). To further characterize this effect, stimuli were grouped into non-social (echolocation) and social(aggression and distress) categories. A linear mixed-effects model revealed that females showed a significantly greater median change in heart rate in response to social stimuli than males (**p < 0.05; Figures 2e**).

**Figure 1:**
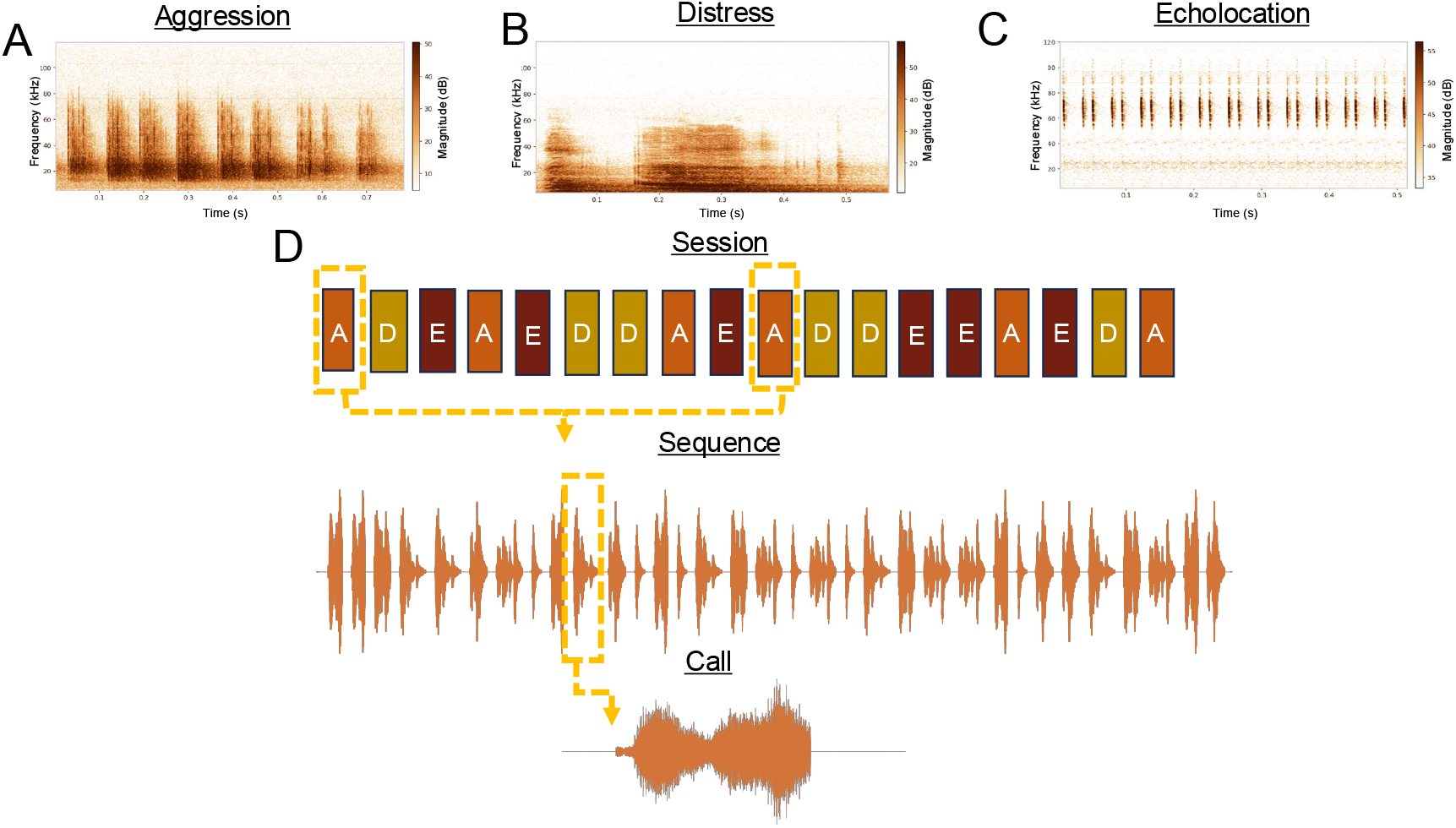
Acoustic Stimuli. (a–c) Spectrograms depicting the energy of each call type over frequency and time, shown in the following order: aggression, distress, and echolocation. **(d)** Schematic of an example recording session during physiological testing, showing organization at the session level, an example stimulus sequence, and an individual call type.

**Figure 2:**
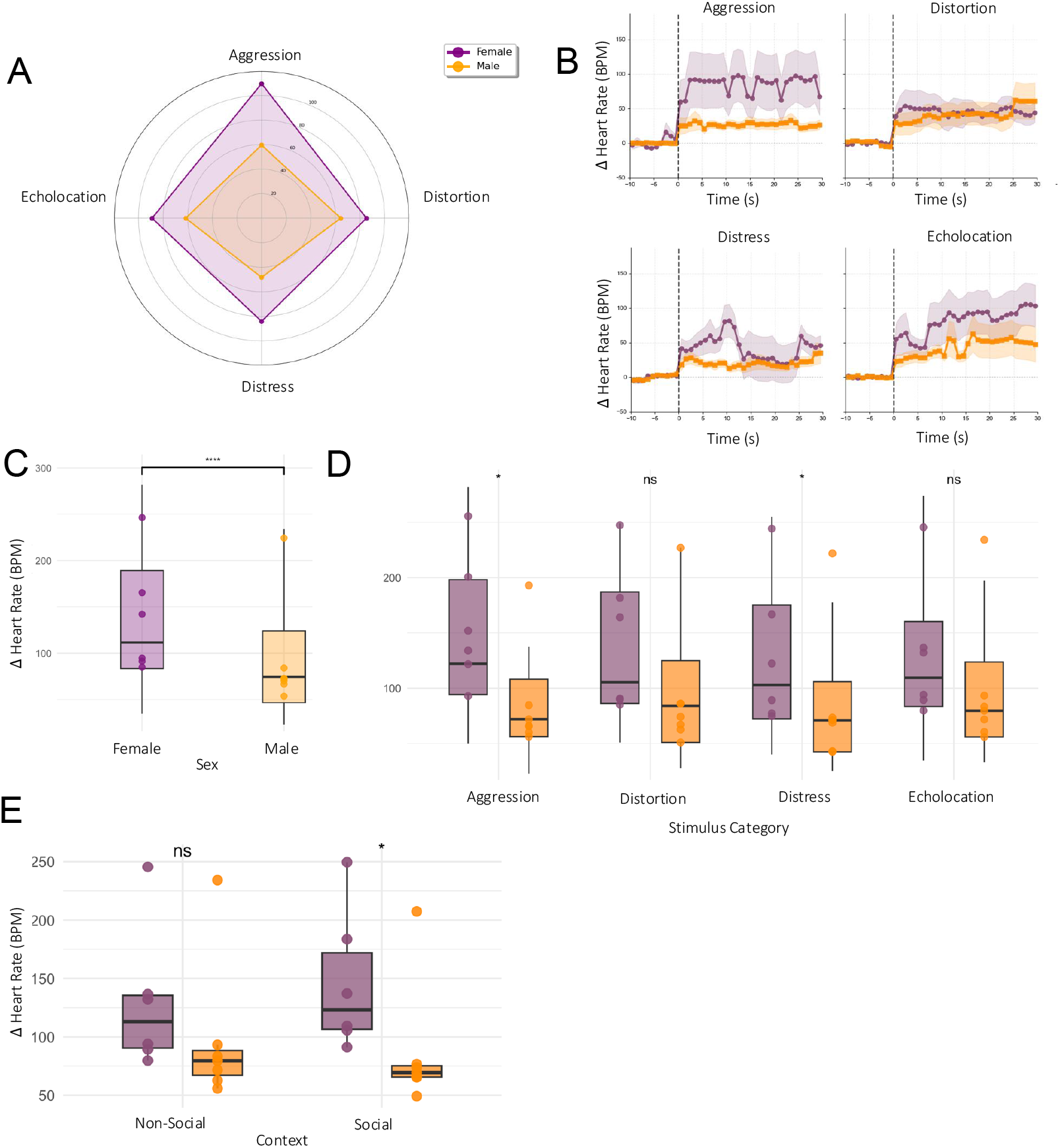
Sex Differences in Heart Rate Responses During the Perception of *R. aegyptiacus* Vocalizations. **(a)** Radar plot depicting the median change in heart rate from baseline for each sex across each stimulus category. **(b)** Average change in heart rate during stimulus presentations for males and females. **(c)** Median change in heart rate between sexes across all stimulus categories (Mann-Whitney U-test, p < 0.001). **(d)** Median change in heart rate between sexes for each stimulus category (Mann-Whitney U-test, p < 0.05). **(e)** Median change in heart rate from baseline in response to social (Aggression, Distress) and non-social (Echolocation) stimuli, separated by sex. Pairwise sex differences within each stimulus category were assessed by the Wilcoxon signed-rank test (*p < 0*.*05, p < 0*.*01*, ** p < 0.001, ns = not significant).

### A1 Contains Call-Selective Units Whose Selectivity is Independent of Frequency Tuning

2,394 MUAs were collected from 6 bats (3 males, 3 females) that underwent electrode implants in the primary auditory cortex, with session recordings conducted during playback of aggression, distress, and echolocation stimuli. Cresyl violet staining was used to verify the electrode implantation site **(Figure 3a)**. When assessing response latency and jitter, social calls (aggression and distress) showed greater variability than the echolocation call, which exhibited a notably short latency **(Figure 3b,3c)**. Tuning analysis of these MUAs revealed no dominant frequency across A1 **(n = 1**,**915; Figure 3d)**, and that tuning did not predict selectivity (Fig, S1). When assessing the selectivity of the MUAs identified via a lifetime sparseness analysis, we found that the majority were selective for distress (4.7%), followed by echolocation (1.9%) and aggression (1.4%), **(Figure 3g)**. Differences were observed in z-scored firing rates between aggression and distress, pointing towards unique firing signatures in response to these call types **(Figure 3h, ***, p<0.001)**. To test whether A1 population activity could discriminate among call categories, we trained a generalized linear model (GLM) decoder on MUA responses using leave-one-out cross-validation. The classifier recovered the preferred category with high accuracy across animals (LOO-CV accuracy: 88.5%, chance: 37.5%, n = 172 units; permutation test, p < 0.001), with per-animal accuracies ranging from 85.7% to 98.4%, demonstrating that the shape of a unit’s firing rate profile across stimulus categories is sufficient to predict its selectivity class (**Figure 3i**).

**Figure 3:**
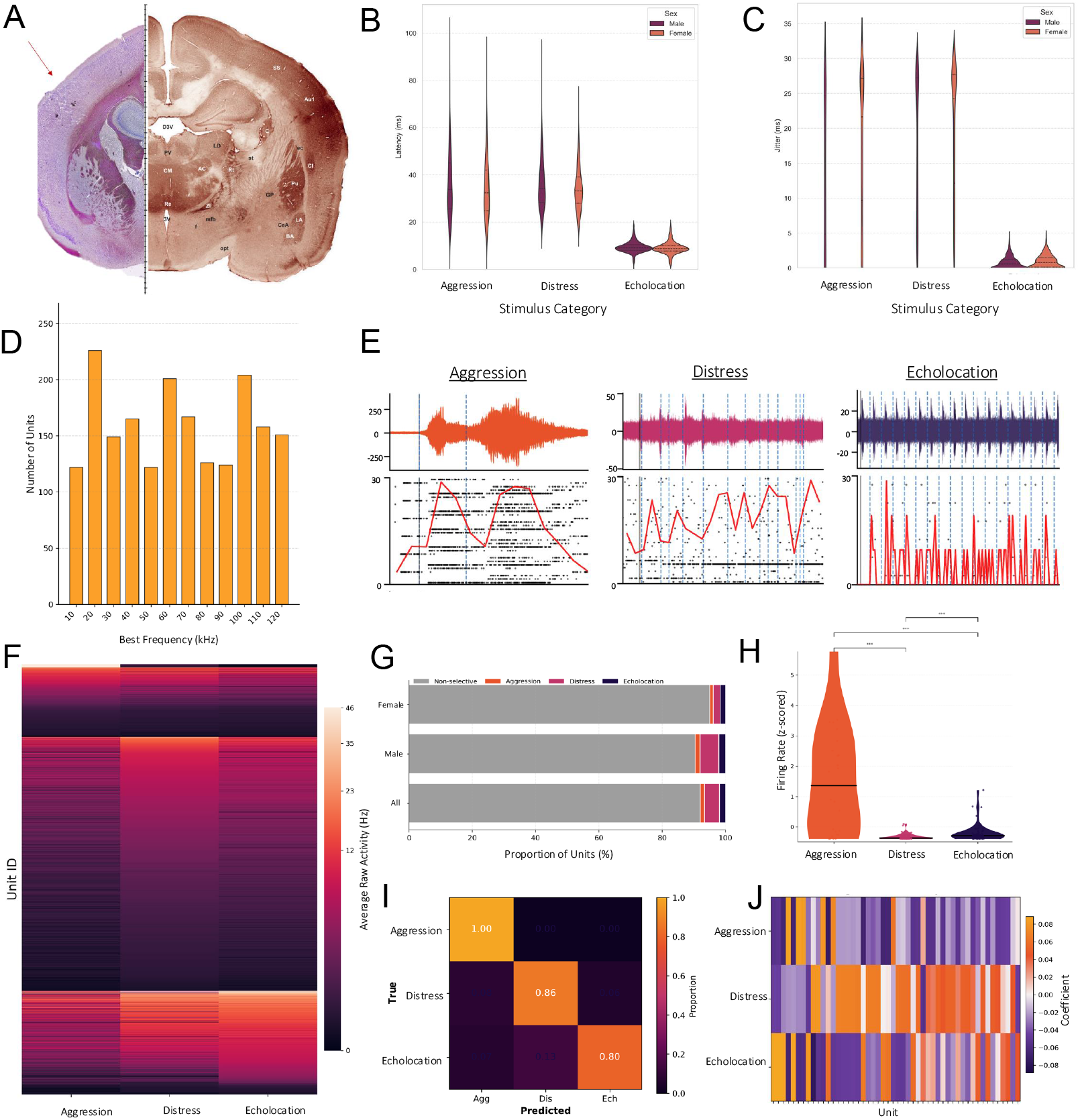
Selectivity within the Auditory Cortex of *R. aegyptiacus* for Different Vocalization Types. **(a)** Histological verification of electrode implant site in the primary auditory cortex. **(b)** Latency values across multiple MUAs by sex. **(c)** Jitter values across multiple MUAs by sex. **(d)** Bar plot quantifying the tuning properties of recorded MUAs. **(e)** Example Raster plots for an MUA across 30 trials of a specific call type (aggression, distress, and echolocation, respectively). **(f)** Heat map representing the average raw activity of MUAs during each stimulus presentation, sorted from highest to lowest activity for each category. **(g)** Bar plot showing the distribution of stimulus selectivity across MUAs as measured by lifetime sparseness (S > 0.5). **(h)** Z-scored firing rate across sound categories. **(i)** Confusion matrix showing LOO-CV decoding performance pooled across all animals. Values indicate the proportion of observations in each true category classified into each predicted category; accuracy = 88.5%; chance = 37.5%. **(j)** Heatmap of fitted GLM coefficients for an example animal, showing the top 50 MUAs ranked by maximum absolute coefficient across categories. Warm colors (orange) indicate positive coefficients (excitatory); cool colors (purple) indicate negative coefficients (inhibitory).

### Approach to Distress Calls Is Gated by Social Context

We assessed approach behavior during distress and white-noise (control) playbacks, both in the presence(social) and absence (nonsocial) of a live conspecific, in 12 adult female bats. Males (n=4) did not exhibit this behavior (Fig. S2). This sample size allowed a within-subject design in which each focal bat received 2 of the 4 stimulus-context conditions, providing 6 bats per stimulus-condition. Bats were randomly assigned conditions for day one and day two trials, ensuring no bat underwent the same stimulus. Bats exhibited the greatest number of approaches during the distress playback under the social context. **(Figure 4b)**. Bats in the distress-social condition spent the most time near the speaker compared to those in the distress non-social condition **(Figure 4c)**. No difference was seen in approach behavior between bats performing on day one or day two, concluding no effect of the order of exposure on approach behavior for each category. **(Figure 4d)**. Across all groups, the distress-social condition produced the closest x-position to the speaker, indicating the greatest proximity. **(Figure 4e)**. To further investigate non-approaching animals during playbacks, micro-behaviors were assessed, including grooming and pointing. The most grooming and pointing was observed during the distress non-social condition. Physical contact with the speaker box occurred exclusively in the distress-social condition **(Figure 4f)**. Together, these results indicate that bats exhibit approach behavior only in the distress social condition, while in the distress non-social condition, they display interest via pointing toward the speaker and anxiety-like behavior, such as grooming.

**Figure 4:**
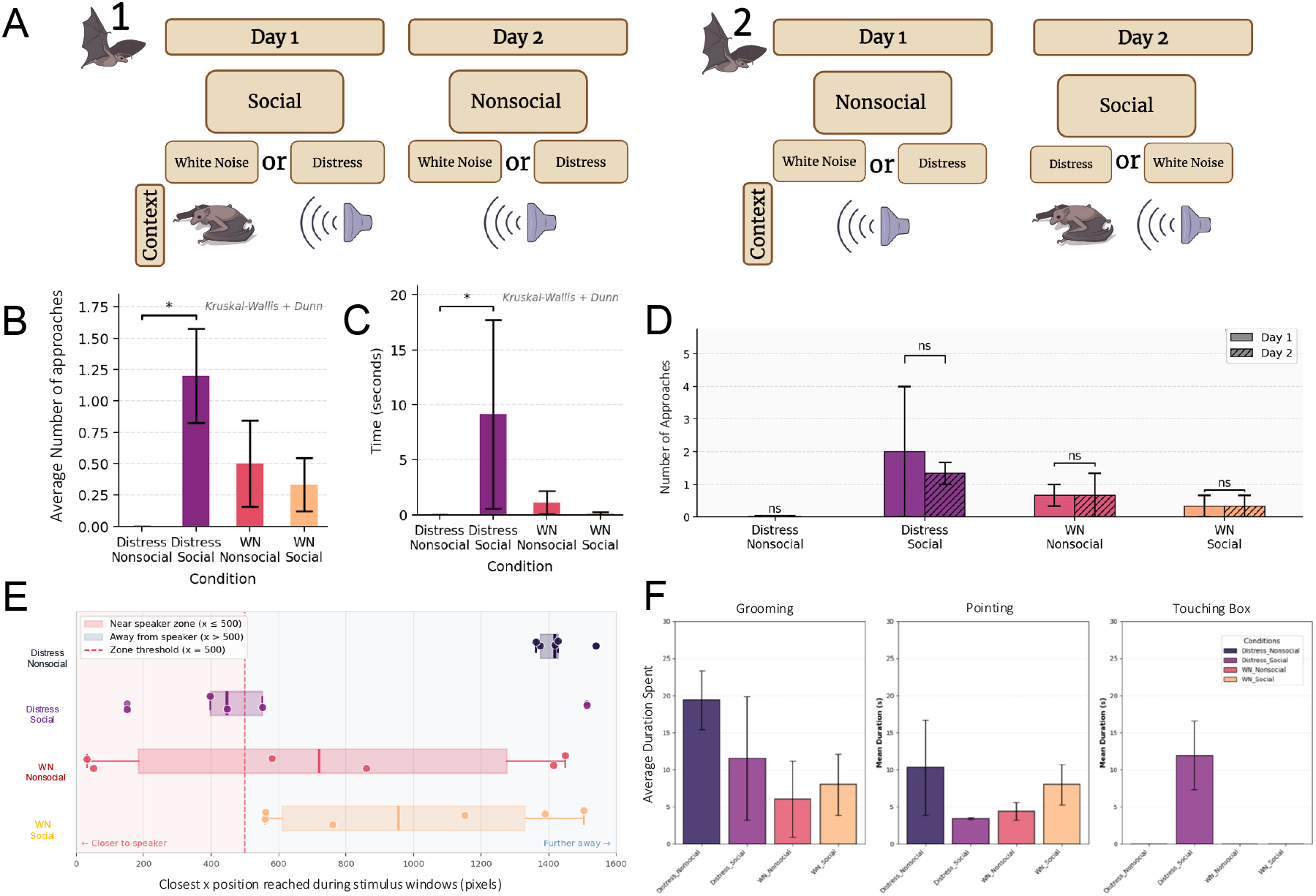
Approach to Distress Calls Is Gated by Social Context. **(a)** Experimental setup for the behavioral paradigm in which two paired bats underwent a playback experiment under different social contexts (Social and Non-Social). A total of 12 female bats were tested, with n = 6 per group. Bar chart depicting the number of approaches toward the speaker for each behavioral context (Distress Non-Social, Distress Social, White Noise Non-Social, White Noise Social). Group differences were assessed using a Kruskal-Wallis test followed by Dunn’s post-hoc test (* p < 0.05, ** p < 0.01, *** p < 0.001). **(c)** Bar chart depicting time spent within the region of interest (ROI), defined as within 200 pixels of the speaker, for each behavioral context (* p < 0.05, ** p < 0.01, *** p < 0.001); statistical comparisons as in (b). **(d)** Bar plots comparing the number of approaches between conditions received on the first day and the second (ns, all p >0.05). Heat maps showing the time spent in each arena for each behavioral condition. **(e)** Horizontal box plot depicting the closest x-position reached by each bat relative to the speaker for each group. **(f)** Box plot depicting the frequency of micro-behaviors assessed via BORIS, including grooming, pointing, and touching the speaker box, averaged across stimulus presentations. BORIS behavioral annotations were conducted by 2 experimenters to ensure unbiased scoring of micro behaviors.

## Discussion

We show that autonomic, cortical, and behavioral responses to social vocalizations in *R. aegyptiacus* are dissociable across contexts. Heart rate and primary auditory cortex responses to social calls are present during passive listening in isolation, whereas approach behavior is not, indicating that social context serves as a necessary gate between sensory encoding and behavior.

### Heart Rate Responses During the Perception of *R. aegyptiacus* Vocalizations are Sex-Dependent

We began by characterizing the physiological valence of these call types in this species, as this has not yet been done. In the literature, it has been shown that *P. abramus* mothers increase heart rate when perceiving pup isolation and echolocation calls [11], and in the wild, both male and female bats approach the speaker when distress calls are played across several species [12-14]. Based on this, we believed we might see an increase in heart rate during passive listening to conspecific vocalizations during the presentations of social communication calls. In assessing heart rate responses, we found that male and female *R. aegyptiacus* exhibit markedly different heart rate responses to conspecific vocalizations during playback trials, with the greatest divergence observed during presentations of social calls rather than nonsocial ones. This distinction points to sex-dependent differences in the intrinsic physiological salience of social vocalizations. Notably, this divergence is not simply a difference in degree change, but also in pattern. Males show a decrease in heart rate from nonsocial to social calls, whereas females show an increase. This allows us to conclude that the two sexes trend in opposite directions. Sex differences in heart rate response peaks suggest that social vocal content is the primary elicitor of dimorphic physiological responses, consistent with broader evidence that gonadal hormones shape autonomic and cortical processing across mammalian brain systems [15].

### A1 Contains Call-Selective Units Whose Selectivity is Independent of Frequency Tuning

After finding sexually dimorphic responses to playbacks of social calls, we asked whether the same pattern would be observed at the neural level. If so, this would indicate selectivity in A1 for social calls. The electrophysiological recordings in A1 provided 2,394 MUAs. Consistent with the acoustic properties of the stimuli, we observed longer latency and jitter values for MUAs responding to social calls, which are longer and more variable in structure, and very short latency and jitter values for responses to echolocation calls, which are short in nature and require precision in timing for navigation. We found that the highest percentage of selective MUAs responded preferentially to distress calls, and that this was most pronounced in males. This was unexpected, as the heart rate data, which showed heightened physiological responses to social calls in females, led us to predict that the female auditory cortex would show the greatest selectivity for these stimuli. This sex-by-stage dissociation suggests that autonomic and cortical measures may index different aspects of social-call processing, with the former perhaps tied to arousal or attentional state and the latter to cortical encoding of acoustic content. Greater male cortical selectivity for distress calls is consistent with a heightened cortical representation of conflict-related signals, although its functional significance remains to be tested in this species. State-dependent modulation of A1 selectivity has been documented in other mammals, where neuromodulators [16] and experience-dependent inhibitory tuning [17] reshape cortical representations of conspecific calls. No single tone frequency dominated across MUAs. Furthermore, we found no significant correlation between best frequency and selectivity across sexes (Fig S1). Having identified distress calls as the most salient stimulus type across both physiological and neural measures, we then asked whether this salience would translate into overt behavioral responses.

### Behavioral Responses to Distress Calls Are Gated by Social Context

Heart rate data demonstrate a physiological emphasis on social calls in females, and the auditory cortical recordings reveal selectivity within A1 for these call types, with the highest percentage of selective MUAs tuned to distress calls. Both datasets were collected in a passive-listening context, prompting the question of whether these patterns would hold in an active-listening context with a live conspecific present. Wild bats have been shown to approach a speaker playing distress calls [18], so we asked whether our captive animals would exhibit similar behavior, whether social context modulates it, or whether call type alone suffices. To address these questions, we designed a playback paradigm that exposed freely flying bats to social/nonsocial conditions across distress and white noise stimuli. We found that approach behavior occurred during distress playbacks in a social context. This condition produced the greatest amount of approach behavior, the most time spent within the ROI, and the closest average proximity to the speaker. Other conditions showed substantially more variability in proximity to the speaker, suggesting generalized curiosity rather than a directed approach. Presentation order had no effect across days, indicating that bats neither habituated nor carried over prior exposure. These results are consistent with the literature, which shows that in another bat species, *S. bilineata*, they exhibit approach behavior toward a speaker emitting distress calls when the emitter is present [19]. Here, conspecific presence alone was sufficient to prompt approach regardless of emitter identity or number of conspecific vocalizations (Fig. S2).

Given the strong physiological signal associated with distress calls, we were curious about the approach behavior being absent in the nonsocial distress condition despite high physiological engagement. Scoring microbehaviors with BORIS [20], we found that bats in this condition showed the most grooming and pointing behaviors, indicating anxiety and attentional direction without approach. This suggests a gating mechanism in which social context is necessary for the approach to be expressed despite physiological and attentional engagement. Notably, this contrasts with contextual-modulation frameworks, in which attention reshapes cortical receptive fields [21]. Here, cortical encoding is preserved, yet approach behavior is absent, suggesting that social context determines whether engagement with a vocalization is expressed as directed action, with implications for how auditory processing and social behavior are coordinated across species. Future work should test whether this gating mechanism generalizes across call types and social configurations, and whether neuromodulators such as oxytocin shape the social-context dependency [22]. Freely-flying recordings in conspecific colonies, building on wireless neural recording approaches in bats [23], would clarify how A1 activity tracks behavior across naturalistic social contexts, and the precise role of the auditory cortex in this gating process.

## Materials and Methods

### Animals

Adult Egyptian Fruit Bats (*Rousettus aegyptiacus*) aged approximately 4 years were housed in a reverse light–dark cycle colony room at controlled temperature and humidity. All procedures were approved by the Institutional Animal Care and Use Committee (IACUC) at the University of Illinois Chicago under protocol 25-027.

### Acoustic Stimuli

Stimuli consisted of calls isolated from recordings made in the colony room and manually annotated. Six exemplars of aggression, distress, and echolocation were taken from the recordings, duplicated, and randomized to produce 30-second bouts of each call type separated by 300 ms of silence in a sequence (**Figure 1**). In addition, for the heart rate monitoring experiment, a distorted stimulus was made by binning the aggression stimuli at 10 ms and randomizing them so that they contained the same spectral features, just not in the same order. Six sequences of each stimulus type were contained in a single recording, separated by 1 minute of white noise. All stimuli were normalized to ≤80 dB SPL at the ear and separated by 1-minute inter-stimulus intervals of white noise. For each experiment, the stimulus content remained the same, but its sequences were rearranged.

### Heart Rate Monitoring

To assess physiological correlates of auditory perception, heart rate was recorded using the MouseOx Plus system [Starr Life Science, Pittsburgh, PA] in awake, passively listening bats.

Twelve adult bats (six females, six males) were habituated to the recording chamber for 1 hour before each session. Each bat was exposed to the full playlist over three days (one trial per day), and heart rate was recorded continuously during stimulus presentation. Changes in heart rate were averaged across trials and analyzed relative to baseline. Baseline heart rate was defined as the lowest heart rate sustained the longest during the trial, to ensure that data collected that was not reflective of baseline heart rate was excluded. The first three presentations of each call were used for analysis, as habituation was observed beyond that point. Data was first analyzed in Python (3.12.13) to produce the radar chart depicting the median change in heart rate across males and females. All other plots were created in R (v2024.04.0).

### Electrophysiological Recordings

To investigate auditory cortical encoding of communication calls, extracellular recordings were performed in awake, passively listening *Rousettus aegyptiacus*. Six bats (three females, three males) were implanted with a 32-channel microelectrode array targeting the auditory cortex (A1). Stereotaxic coordinates were adjusted individually based on cranial width and length due to inter-individual anatomical variability. Surgical procedures followed protocols approved by the IACUC and included post-operative recovery for 24 hours before recording. Recordings were conducted over 2 weeks. Auditory stimuli included 30-second randomized bouts of distress, aggression, echolocation, and white noise. Each bout was composed of exemplar calls annotated and verified by spectrogram shape, frequency, and intensity. Frequency tuning curves (10–120 kHz, 80 dB SPL, tapered on/off ramps) were presented to determine the best frequency at each site. Electrode placement was confirmed via cresyl violet staining.

### Speaker Playback Test

To assess the behavioral salience of vocalizations, 12 adult female and 4 adult male focal bats were exposed to playbacks of distress vocalizations or to our white-noise control on 2 days separated by 1 week. On Day 1, each bat was assigned to either a social context (with a live conspecific present) or a nonsocial context. They were also assigned a stimulus: either distress or white noise. After the first testing day, we allowed 1 week to pass to prevent habituation to the task and the speaker. Each bat was then presented with the opposing combination of Day 1. If they received distress under a social condition on Day 1, then they received white noise under a nonsocial condition on experimental Day 2. These two stimuli were played via an Avisoft speaker placed at the center of the chamber. Stimuli included 30-second exemplars of distress and white noise, matched in intensity (≤80 dB SPL) and separated by inter-stimulus intervals (1 min). All tests were conducted during the bats’ active night phase. Audio annotations and spectrogram validation were performed using Audacity [24] and Kaleidoscope [25]. BORIS was used to quantify micro-behaviors such as grooming, pointing, and speaker contact, which were further analyzed in Python. Calls recorded during trials were manually annotated. Analysis was performed in R.

### Data Analysis for Electrophysiology

Data collected during session recordings were spike sorted using Kilosort4 [26] with default parameters. Sorted MUAs were manually curated by visual inspection of waveform shape, inter-spike interval distributions, and cluster separation. SpikeInterface (Python 3.12.13) [27] was used for data handling, preprocessing, and visualization of spike waveforms and firing activity. Lifetime sparseness was computed in Python using the standard formulation for lifetime sparseness [28]. MUAs with sparseness values greater than 0.5 were classified as sparse. To assess whether the preferred stimulus category could be predicted from unit firing profiles, Claude (Anthropic) was used to fit a generalized linear model (L2 regularization, C = 0.1, balanced class weights) per animal using leave-one-out cross-validation (LOO-CV) across selective units. Each unit was represented by four features (z-scored) per stimulus category: average firing rate, response standard deviation, peak firing rate, and a pairwise modulation index for each of the three categories. Ground-truth preferred category was assigned using the sparseness threshold. Predicted labels were pooled across animals for summary accuracy and confusion matrix computation. Permutation testing (n = 1,000 shuffles of preferred-category labels within animal) generated a null distribution against observed accuracy.

### Data Analysis for Behavioral Experiment

Behavioral activity was quantified using a custom Python computer vision pipeline. Video recordings were acquired using a fixed overhead camera (1600 × 1000 pixels) positioned above the behavioral arena. Animals were detected using a deep learning-based object detection model implemented with YOLOv8, trained on 800 manually annotated frames spanning variation in lighting, posture, and background structure. The model was iteratively retrained to account for challenging lighting conditions and background artifacts. Inference was performed using a fixed confidence threshold of 0.25, and a minimum displacement threshold of 1 cm per frame was applied to reduce contributions from minor positional jitter.

## Acknowledgments

This work was supported by a Diversity Supplement from the National Institute on Deafness and Other Communication Disorders (NIDCD), and an R00 DC019145. The author would like to acknowledge the use of Claude (Anthropic) in the editing and data analysis process.

