## supplemental information for "Passive Physiological Responses Fail to Predict Context-Dependent Action Selection in Bats"

**This PDF file includes:**

Supporting text
Figures S1 to S2 (BF information, and then vocalizations?)
Legends for Datasets S1 to Sx

### Supporting Information Text

### Figures

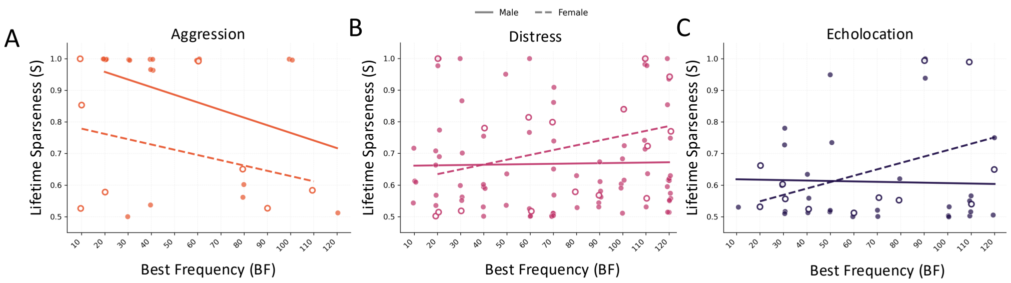


**Fig. S1.** Sex-specific analyses of lifetime sparseness and Best Frequency. Scatter plots of lifetime sparseness (S) as a function of best frequency (BF, kHz) for male (filled circles, solid regression line) and female (open circles, dashed regression line) MUAs in three vocalization categories: **(A)** aggression-selective, **(B)** distress-selective, and **(C)** echolocation-selective. Linear regression statistics are as follows — Aggression: males $R^2 = 0.136$, $p = 0.101$, $n = 21$; females $R^2 = 0.115$, $p = 0.412$, $n = 8$. Distress: males $R^2 = 0.0005$, $p = 0.838$, $n = 81$; females $R^2 = 0.102$, $p = 0.228$, $n = 16$. Echolocation: males $R^2 = 0.0009$,$p = 0.878$, $n = 28$; females $R^2 = 0.192$, $p = 0.154$, $n = 12$. No significant relationship was detected for either sex in any category (all $p > 0.05$).

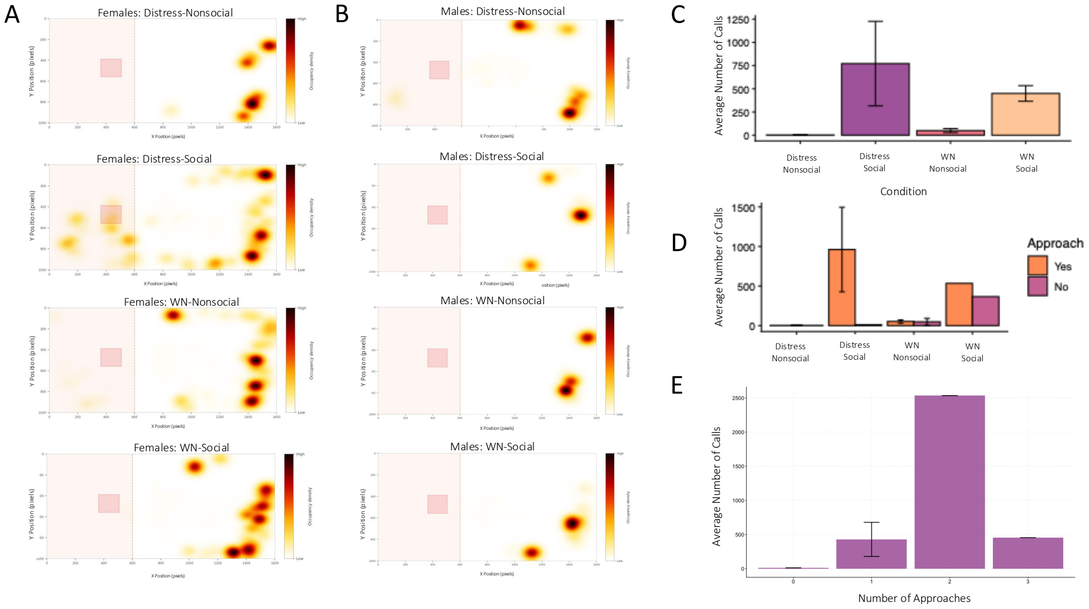


**Fig. S2. (A–B)** Trajectory-averaged heat maps of zone occupancy density for females **(A)** and males **(B)** during stimulus presentation. Males showed no approach behavior during stimulus presentations across all conditions (n = 4; 1 per condition). **(C)** Average number of vocalizations produced across the four conditions (Distress Nonsocial, Distress Social, White Noise Nonsocial, White Noise Social) in females (n = 12, 6 per group). No significant differences in vocalization number were observed across conditions (all $p > 0.05$). **(D)** Average number of vocalizations per condition separated by whether approach behavior was displayed (Yes/No). No consistent relationship between vocalization number and approach behavior was observed across conditions, though small sample sizes limit interpretation. **(E)** Average number of vocalizations as a function of number of approaches during stimulus presentation in females (n = 12). No clear monotonic relationship between vocalization number and approach frequency was observed. These findings suggest vocalization number alone may not predict approach behavior in females.
